# Mapping Sleep’s Oscillatory Events as a Biomarker of Alzheimer’s Disease

**DOI:** 10.1101/2023.02.15.528725

**Authors:** Rachelle L. Pulver, Eugene Kronberg, Lindsey M. Medenblik, Vitaly O. Kheyfets, Alberto R. Ramos, David M. Holtzman, John C. Morris, Cristina D. Toedebusch, Stefan H. Sillau, Brianne M. Bettcher, Brendan P. Lucey, Brice V. McConnell

## Abstract

**Objective:** Memory-associated neural circuits produce oscillatory events within single-channel sleep electroencephalography (EEG), including theta bursts (TBs), sleep spindles (SPs) and multiple subtypes of slow waves (SWs). Changes in the temporal “coupling” of these events are proposed to serve as a biomarker for early stages of Alzheimer’s disease (AD) pathogenesis.

**Methods:** We analyzed data from 205 aging adults, including single-channel sleep EEG, cerebrospinal fluid (CSF) AD-associated biomarkers, and Clinical Dementia Rating® (CDR®) scale. Individual SW events were sorted into high and low transition frequencies (TF) subtypes. We utilized time-frequency spectrogram locations within sleep EEG to “map” the precision of SW-TB and SW-SP neural circuit coupling in relation to amyloid positivity (by CSF Aβ_42_/Aβ_40_ threshold), cognitive impairment (by CDR), and CSF levels of AD-associated biomarkers.

**Results:** Cognitive impairment was associated with lower TB spectral power in both high and low TF SW-TB coupling (p<0.001, p=0.001). Cognitively unimpaired, amyloid positive aging adults demonstrated lower precision of the neural circuits propagating high TF SW-TB (p<0.05) and low TF SW-SP (p<0.005) event coupling, compared to cognitively unimpaired amyloid negative individuals. Biomarker correlations were significant for high TF SW-TB coupling with CSF Aβ_42_/Aβ_40_ (p=0.005), phosphorylated-tau_181_ (p<0.005), and total-tau (p<0.05). Low TF SW-SP coupling was also correlated with CSF Aβ_42_/Aβ_40_ (p<0.01).

**Interpretation:** Loss of integrity in neural circuits underlying sleep-dependent memory processing can be measured for both SW-TB and SW-SP coupling in spectral time-frequency space. Breakdown of sleep’s memory circuit integrity is associated with amyloid positivity, higher levels of AD-associated pathology, and cognitive impairment.

## Introduction

Sleep dysfunction is hypothesized to share a bidirectional relationship with Alzheimer’s disease (AD) pathology^1,2^, and there is growing interest in understanding the neurophysiological properties of sleep that are most strongly associated with neurodegeneration. Among many putative neuroprotective attributes of sleep, slow wave activity (SWA) stands out for both the robust data supporting plausible neuroprotective mechanisms^2,3^, and the readily quantified SWA metrics that can be obtained from widely available single-channel electroencephalography (EEG)^4-6^. Loss of SWA correlates with age^7-11^ and neurodegenerative processes including Alzheimer’s and Parkinson’s disease^2,12^. Further, in preclinical AD, SWA loss occurs in association with amyloid deposition rates^13^, and the presence of tau pathology^4^.

There are extensive data supporting SWA’s role in synaptic homoeostasis^14^, and regulation of synaptic plasticity is thought to support SWA’s role in sleep-dependent memory consolidation^15,16^. Within SWA, multiple types of oscillatory events occur in association with replay of memory sequences, mirroring wake-like experiences in the patterns of neuronal activity^17-23^. Oscillatory components of SWA’s memory playback include slow waves (SWs), theta bursts (TBs), and sleep spindles (SPs), and together they form nested, or “coupled”, complexes with one another during SWA^24-29^. Experiments in preclinical models have further demonstrated that memory processing can be disrupted or enhanced via specific modulation of SW and SP events^30-32^.

Timing irregularities of SW and SP coupling have been correlated with amyloid^33^ and tau^34^ in positron emission tomography (PET) imaging studies. Further, subpopulations of SW and SP oscillatory events demonstrate unique properties, and distinct relationships exist within SW events defined by either high versus low transition frequencies in the context of aging^35^, amyloid positivity^33^, and cognitive processes^33,36^. Theta burst events are detectable prior to the troughs of SWs^5,6,25,26,37^ and play a role in normal sleep-dependent memory processing^37,38^, although their relationships to SW subpopulations, and their potential changes in aging and neurodegenerative processes, have yet to be formally assessed.

Remarkably, a simple single channel of EEG recording is sufficient to probe the integrity of memory-associated oscillatory events, thus opening the door to deploy inexpensive “wearable” devices in the home setting for monitoring brain health and early signs of neurodegenerative disease. Nonetheless, translation of this technology into clinical application will require significant additional work, including steps to better characterize the various subtypes of SWA-associated oscillatory events in both normal and pathological processes. Here we sought to make such advancements by innovating signal processing methods to map the spectral coordinates of individual oscillatory events in time-frequency space, thus providing a metric of both temporal and frequency precision of the neural circuits underlying sleep-dependent memory consolidation.

In this study, we utilized this novel time-frequency spatial mapping to examine the properties of several key oscillatory events (including high and low transition frequency SW subtypes, SPs, and TBs) within a large and well-characterized cohort of older adults. We hypothesized that the time-frequency precision and event-specific EEG power of SW-coupled oscillations would correlate with amyloid positivity and cognitive impairment. We further assessed correlations between metrics of oscillatory event coupling and cerebrospinal fluid (CSF) levels of core AD biomarkers.

## Methods

### Participant Sample

Community-dwelling participants from a longitudinal cohort at the Knight Alzheimer Disease Research Center (ADRC) at Washington University in St. Louis were selected for analysis (n=205). All data were collected with written informed consent under research protocols approved by the Washington University in St. Louis Institutional Review Board. Participants were selected for this study if they had completed at least 3 nights of single-channel EEG recording, 1 night of monitoring with a home sleep apnea test, genotyping for APOE4 status, one lumbar puncture for Alzheimer’s biomarkers within 1 years of sleep recordings, anda Clinical Dementia Rating® (CDR®)^39^ within 2 years of sleep recordings. All participants were either cognitively unimpaired (CDR 0) or very mildly cognitively impaired (CDR 0.5) with the exception of one participant who was mildly cognitively impaired (CDR1).

### EEG and Apnea Data Acquisition

Overnight EEG recordings were acquired as previously described^40^. Briefly, longitudinal EEG recordings were obtained from participants at home up to 6 nights using a single-channel EEG device worn on the forehead (with sensors at approximately AF7, AF8, and Fpz) and with a sampling rate of 256 samples per second (Sleep Profiler, Advanced Brain Monitoring). Resulting EEG were visually scored by registered polysomnographic technologists using criteria adapted from the American Academy of Sleep Medicine guidelines^41^. An additional a one-night home sleep apnea test was utilized as previously described^36^ to measure hypopneas greater than 4% oxygen desaturation criteria and compute an apnea–hypopnea index (AHI) for each participant (HSAT; Alice PDx, Philips Respironics Inc, Murrysville, PA).

### Slow Wave Detection

Raw EEG timeseries data was processed with MATLAB R2021b (MathWorks, Inc., Natick, MA, USA.). Slow waves were identified via automated zero-crossing detection as previously described^6^. Briefly, slow wave detection was performed from forehead electrodes, roughly FP1-FP2 montage, from sleep stages N2 and N3. Epochs with un-scorable data were excluded from analysis. Automated management of high amplitude artifacts was accomplished via exclusion of EEG segments exceeding 900 µV after detrending data with sliding window of three seconds across raw data. A high amplitude, repeating artifact from the recording device was also excluded by thresholding. Discrete Fourier transform (DTF) was computed using a fast Fourier transform (FFT) algorithm for the affected frequency region (15hz - 17hz) and artifact-containing regions were removed if the DFT vector values exceeded 8 µV. EEG data was subsequently detrended and band-pass filtered in a forward and backward direction using a 6th-order Butterworth filter between 0.16-4 Hz. Zero crossings were identified to detect negative and positive half-waves, and slow wave events were identified when the half-wave pairs approximated a frequency range of 0.4 to 4 Hz. Minimal and maximal half-wave amplitudes were measured, and slow waves with both positive and negative maximum amplitudes in the top 50% of all waves were selected for subsequent coupling analysis. An upper threshold of +/- 200 µV for zero crossing pairs was utilized to reduce misidentification of non-slow wave events. A further reduction of false identifications was accomplished by rejecting all zero crossing pairs with peak/trough amplitudes exceeding four standard deviations from the mean min/max zero crossing pair values for each subject.

### Spindle and Theta Burst Identification

Spindle and theta burst event identification was performed using established methods.^5,6^ Briefly, EEG data was detrended and bandpass filtered in a forward and backward direction using a 3rd-order Butterworth filter between 10-13.5 Hz for late-fast spindles and 4-8 Hz for theta bursts. Note that early-fast spindles are more prominent in central recording locations^5^ and were not consistently detected in the FP1-FP2 channel, and therefore were excluded from analysis. Maximum spindle envelopes were calculated and an amplitude threshold of 75% percentile of the root mean squared value with a length window of 0.5 sec to 3.0 sec was used to define spindle events and theta bursts. An absolute threshold of 40 µV in range was used to eliminate artifacts and only spindle/theta burst envelopes within 8 standard deviations from the mean amplitude values were selected for analysis.

### Separation of Slow Wave by Transition Frequency

Slow wave events were categorized as high versus low transition frequency in accordance with previously published methods^33,35^. The distance between the trough and peak of each slow wave was calculated and resultant halfwave was converted into a frequency value in Hertz. A cutoff of 1.2 Hz was then used to separate all detected slow waves into two populations, as previously described (Figure 1)^33,35^.

**Figure 1:**
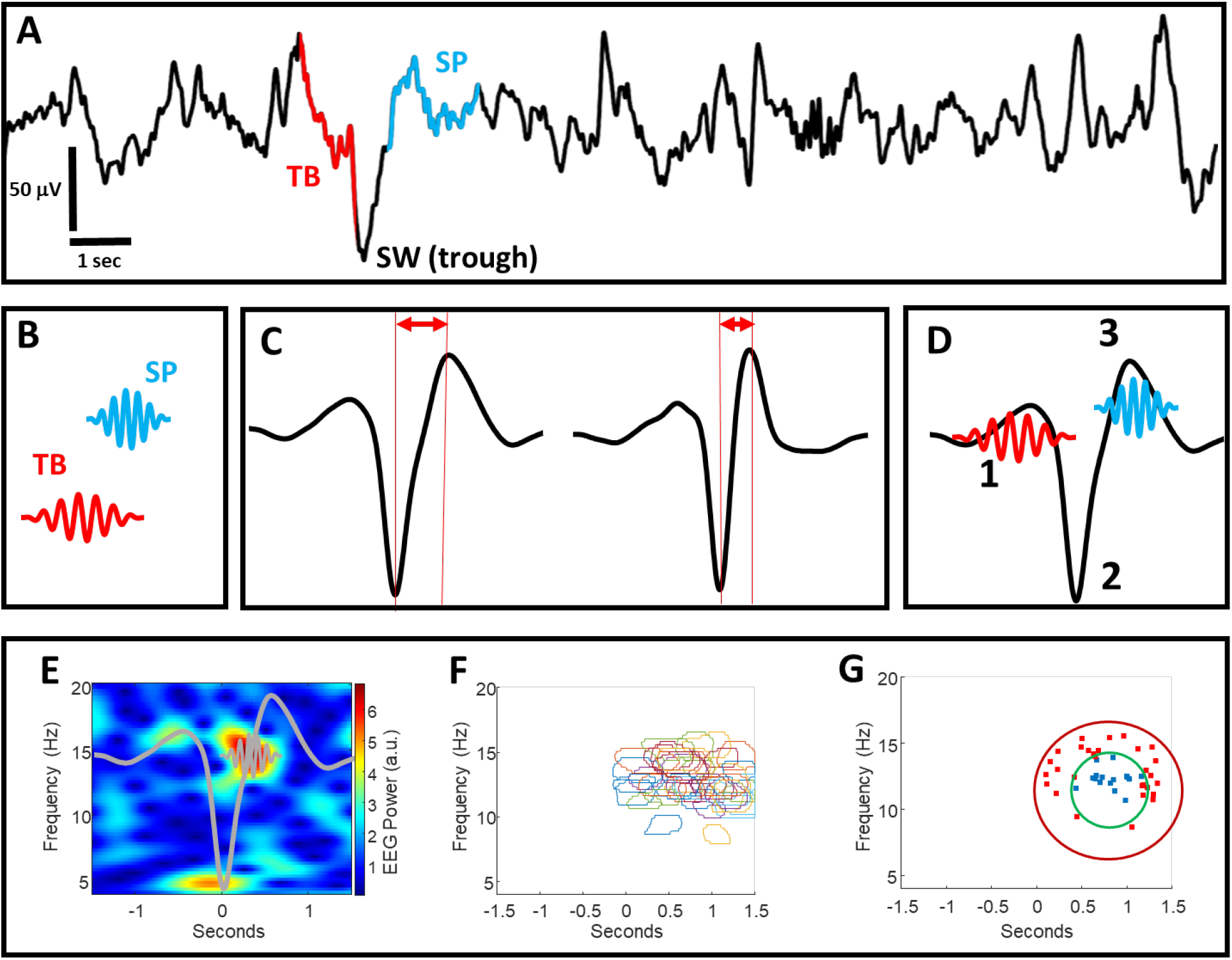
A) An illustrative example of raw sleep EEG data containing an identified theta burst (TB) event and a spindle (SP) event, each temporally coupled to the trough of a slow wave (SW) event. Note that not all SW events can be matched to coupled TB or SP events. B) A conceptual illustration of TB and SP oscillatory events that are individually identified from raw EEG (illustration not to scale). C) Slow wave (SW) oscillations are identified from raw EEG and characterized as having low or high transition frequencies by measuring the distance from their troughs to peaks. D) Temporal sequences of “1” TBs, “2” SWs, and “3” SPs create “coupling” as the events co-occur in fixed time windows from one another (illustration not to scale). E) An example of a normalized time-frequency EEG spectrogram containing an individual SP event coupled to the trough of an individual SW (SW and SP shapes are superimposed as a conceptual illustration, not to scale). F) An example of event detection wherein the time-frequency location of individual SP events from multiple spectrograms are outlined and superimposed. G) An example of precision event topographical localization. The precision of SP event centroids in panel “F” is calculated by measuring the number of SP event centroids within the inner ring, divided by the total number within the outer and inner ring areas. Abbreviations: SP = spindle, SW = slow wave, TB = theta bursts.

### Detection of Slow Wave Coupling with Spindles and Theta Bursts

Individual SW events were sorted by their co-localization with SPs and TBs in the time domain as previously described^5,6^. Spindles occurring within 0 sec to 1.5 sec from the trough of each SW were classified as a coupled event. Theta bursts occurring within - 0.5 sec to 0.2 sec from the trough of each slow wave were classified as a coupled event.

### Time-Frequency Spectrogram Analysis

Time-frequency wavelet spectrograms of slow wave-coupled spindles and theta bursts were created via established methods.^5,6^ Briefly, troughs of each slow wave were centered in 5-second intervals of EEG data and matched to 5-second baseline intervals immediately preceding slow wave events (excluding baseline segments containing slow wave events). A Morlet-wavelet transformation (65 cycles from 4 Hz to 10 Hz) was applied to the unfiltered EEG for slow wave and baseline segments between 4 Hz and 20 Hz in steps of 0.25 Hz with varying wave numbers (65 cycles from 4 Hz to 10Hz with a step size of 0.0938 to match the frequency step size). The mean of baseline regions was used to normalize the amplitude of the mean Morlet-wavelet transformation of all 5-second slow wave-adjacent regions. Time-frequency (TF) windows were defined within time-frequency spectral space for quantification of baseline-normalized EEG power. Theta bursts were defined by a TF-window between 4 Hz and 6.5 Hz at -0.5 s to 0.2 s from the average slow wave trough. Spindle TF-window normalized EEG power was measured between 10 Hz and 13 Hz at 0.3 s to 1.3 s from the average slow wave troughs.

### Precise Event Topographical Localization Analysis

Time-frequency wavelet spectrograms of normalized EEG power were utilized to perform precise event topographical localization (PETL) to determine the location of each TB and SP event in time-frequency space. Briefly, the individual spectrogram images for each SW-TB and SW-SP event were processed via the MATLAB image processing toolbox to detect oscillatory events within a time-frequency space of 10-18 Hz and 0-1.4 sec from the SW trough. Each spectrogram was converted into a binary image using the ‘imbinarize’ MATLAB function, where pixels above an intensity threshold of 0.65 were set to 1 and all other pixels were set to zero. Subsequently, centroids of round-shaped oscillatory events (each exceeding >0.4 roundness factor; >1000 pixels/499,850 total pixels) were mapped within two circular target zones with radii spanning 4 Hz and 2 Hz in frequency space, respectively. A precision metric was calculated by dividing the number of event centroids within the inner circular target zone by the total number of event centroids in the inner and outer circular target zones (expressed as a percentage of events within the inner circle target zone; Figure 1)

### CSF Biomarker Acquisition and Thresholding

CSF collection was performed as previously described in a standardized protocol^42^. Briefly, lumbar punctures were performed at 8:00 a.m. with a 22-gauge Sprotte spinal needle and aliquoted into polypropylene tubes after low speed centrifugation. Samples were stored at -80^°^C until analysis. Concentrations of amyloid-β42, amyloid-β40, t-tau, and p-tau 181 were obtained using previously described protocols via automated electrochemiluminescence immunoassay (LUMIPULSE G 1200, Fujirebio), and thresholding for amyloid positivity was performed as previously described^43,44^.

### Statistical Analysis

Statistical analysis was performed with SAS v9.4 (SAS Institute Inc., Cary, NC, USA). Demographics and other subject level variables were compared among groups, using ANOVA type models for continuous or scale variables, and chi-square/Fisher’s exact association test for categorical variables. Logarithmic transforms were considered for right skewed distributions. Negative binomial count rate models, with robust standard errors, were considered for counts of sleep events.

EEG variables were logarithmically transformed and analyzed with mixed models to compare cognitive groups, adjusted for age, sex, years of education, APOE 4 (yes vs no), and AHI. A random intercept was invoked for repeated measures on a subject across multiple nights. Different residual variances were allowed for different treatment. An omnibus F test for performed for differences among the cognitive groups, followed by pair-wise comparisons with the Tukey-Kramer adjustment. Additive differences on the logarithmic scale were back transformed into ratios and percent differences on the original scale. Estimates, 95% confidence intervals, and p-values for testing the null hypothesis of no difference were reported.

Partial Spearman correlations were run for the relationships between EEG variables (subject averaged on the logarithmic scale) and biomarkers, adjusted for age, sex, years of education, APOE 4 (yes vs no), and AHI. The p-values for the hypothesis test of no correlation were obtained, and confidence intervals were calculated using the Fisher Z transform. Sample correlations, 95% confidence intervals, and p values were reported. A Benjamini-Hochberg procedure was used to control the false discovery rate (FDR) among biomarker correlation with the EEG precision variables for the different spectra and select the ones which remained statistically significant. Two-sided alpha = 0.05.

## Results

A total of 205 participants met criteria for analysis. Subdividing the participants by amyloid positivity and CDR status resulted in 105 participants who were cognitively normal by CDR testing and amyloid negative by Aβ42/40 CSF cutoff (herein referred to as the Aβ-CU group). An additional 69 participants were cognitively normal by CDR testing and amyloid positive by CSF cutoff (herein referred to as the Aβ+CU group), and 31 participants were cognitively impaired by CDR testing and amyloid positive by CSF cutoff (herein referred to as the Aβ+CI group). Demographics and sleep study metrics are provided in Table 1. Age was not significantly different between Aβ-CU and Aβ+CI individuals, although the Aβ+CU individuals were on average ∼2 years older than Aβ-CU individuals (p<0.05). There were no statistically significant differences in education, sex, or AHI between the groups. The APOE4 allele demonstrated an expected higher prevalence among Aβ+CU and Aβ+CI adults, compared to Aβ-CU adults (p<0.001). Sleep staging metrics were similar across groups, with no significant differences observed (Table 1).

**Table 1.** Demographics and Staging. Abbreviations: Aβ = amyloid; CU = cognitively unimpaired; CI = cognitively impaired; TST= total sleep time; REM = rapid eye movement; SOL= sleep onset latency; SE = sleep efficiency; WASO = wake after sleep onset ^a^ p < 0.05 Aβ− CU vs Aβ+ CU ^b^ p < 0.001 Aβ− CU vs Aβ+ CU ^c^ p < 0.001 Aβ− CU vs Aβ+ CI

| **Demographics** | **Aβ- CU** | **Aβ+ CU** | **Aβ+ CI** |
| --- | --- | --- | --- |
| No. participants (%) | 105 (51) | 69 (34) | 31 (15) |
| Age at sleep study, years, median (IQR) | 71 (69 – 75) | 73 (71 – 79)<sup>a</sup> | 74 (69 – 79) |
| Males, *n* (%) | 46 (43.8) | 30 (43.48) | 20 (64.52) |
| Education, years, median (IQR) | 16 (15 – 18) | 17 (15 – 18) | 18 (15 – 19) |
| Number of APOE4-positive (%) | 21 (20) | 35 (50.72)<sup>b</sup> | 21 (67.74)<sup>c</sup> |
| Aβ<sub>42</sub>/Aβ<sub>40</sub> ratio, median (IQR) | 0.092 (0.087 – 0.097) | 0.048 (0.041 – 0.059)<sup>b</sup> | 0.044 (0.039 – 0.052)<sup>c</sup> |
| **Sleep Staging** |  |  |  |
| Minutes, median (IQR) |  |  |  |
| TST | 371.6 (340.38-399.5) | 374.75 (331.60 – 408.92) | 401.83 (352.28 – 446.20) |
| Stage N1 | 29.67 (23.25 – 37.6) | 31.08 (22.33 – 38.0) | 26.70 (20.92 – 40.00) |
| Stage N2 | 254.33 (215.92 – 284.5) | 258.5 (226.67 – 289.30) | 271.4 (218.58 – 318.05) |
| Stage N3 | 2.13 (0.33 – 9.3) | 2.58 (0.3 – 12.58) | 1.17 (0.11 – 8.65) |
| REM | 83.83 (60.63 – 96.58) | 77.6 (63.20 – 88.38) | 86 (63.29 – 106.17) |
| REM latency | 86.17 (62.33 – 123.80) | 77.67 (56.33 – 114.33) | 76 (58.83 – 136.73) |
| SOL | 18 (10.50 – 24) | 12.67 (7.75 – 24.20) | 14.8 (6.74 – 21.33) |
| SE | 81.14 (73.54 – 85.25) | 80.53 (75.82 – 84.84) | 81.37 (76.29 – 87.78) |
| WASO | 67.70 (47 – 105.10) | 74.6 (50.58 – 90.83) | 74.1 (39.65 – 99.27) |
| N1 % | 7.99 (6.34 – 10.91) | 8.36 (6.12 – 10.35) | 7.71 (5.72 – 9.71) |
| N2 % | 67.70 (63.65 – 72.10) | 68.01 (62.27 – 75.10) | 67.59 (63.70 – 75.86) |
| N3 % | 0.61 (0.10 – 2.16) | 0.66 (0.09 – 3.01) | 0.35 (0.03 – 2.21) |
| REM % | 22.50 (18.33 – 25.25) | 20.31 (16.38 – 25.18) | 21.13 (16.83 – 27.42) |

### Detection of Memory-Relevant Events from Sleep EEG

We observed similar numbers of overall SW, TB, and SP events between Aβ-CU, Aβ+CU, and Aβ+CI adults (Table 2). Slow wave events were sorted into subtypes of high and low transition frequency at 1.2 Hz cutoff, and no significant differences in event counts were identified between groups. Temporal coupling of SWs to both SPs and TBs was also comparable between groups, without any significant differences identified (Table 2).

**Table 2.** Sleep Event Counts. Number of detected events are presented as median (IQR). There were no significant differences between groups. Abbreviations: Aβ = amyloid; CU = cognitively unimpaired; CI = cognitively impaired; IQR= interquartile range; SP = spindle; SW = slow wave; TB = theta bursts

| **Median (IQR)** | **Aβ- CU (n=105)** | **Aβ+ CU (n=69)** | **Aβ+ CI (n=31)** |
| --- | --- | --- | --- |
| Total SWs | 15,138<br>(10,188 – 20,550) | 15,763<br>(11,027 – 20,806) | 13,856<br>(11,208.50 – 21,228.50) |
| Total SWs per night | 2,549.50<br>(1,698 – 3,454.17) | 2,870.25<br>(1,985.5 – 3,467.67) | 2,631.50<br>(1,980.45 – 3,538.08) |
| High transition frequency SWs | 5,252<br>(3,237.50 – 7,151.50) | 6,048<br>(3,850 – 8922) | 5,405<br>(3,569 – 7,559) |
| High transition frequency SWs per night | 917<br>(544.20 – 1,201.33) | 1,055.25<br>(683.83 – 1,490.83) | 1,063.33<br>(697.90 – 1,320.92) |
| Low transition frequency SWs | 9,414<br>(6,312.50 – 12,050.50) | 10,642<br>(7,077 – 11,581) | 8,124<br>(6,553 – 13,747) |
| Low transition frequency SWs per night | 1,588<br>(1,059.17 – 2,015.33) | 1,797.50<br>(1,277 – 2,087.67) | 1,636.83<br>(1,209.67 – 2,315.65) |
| Total TBs | 20,946<br>(15,426 – 26,058.50) | 22,646<br>(17,378 – 26,532) | 20,504<br>(17,529.50 – 26,729) |
| TBs per night | 3,579.83<br>(2,566.17 – 4,360) | 3,997.33<br>(2,971.75 – 4,622.50) | 3,683.60<br>(3,271.42 – 4,657.22) |
| Total SPs | 23,016<br>(17,263.50 – 28,177.50) | 24,963<br>(18,439 – 28,942) | 22,391<br>(17,864 – 29,271.50) |
| SPs per night | 3,836.83<br>(2,894 – 4,759.33) | 4,362.83<br>(3,221.17 – 4,909.83) | 4,257.50<br>(3,250.67 – 4,946.17) |
| High transition frequency SW-TB coupling | 1,624<br>(1,021.50 – 2,301) | 1,810<br>(1,194 – 2,742) | 1,673<br>(1,143.50 – 2,245) |
| High transition frequency SW-TB coupling, per night | 270.67<br>(160.33 – 388) | 323<br>(205.67 – 457) | 304<br>(212.40 – 394.17) |
| High transition frequency SW-SP coupling | 2,623<br>(1,582.50 – 3,466.50) | 3,107<br>(1,948 – 4,298) | 2,694<br>(1,766 – 3,750.50) |
| High transition frequency SW-SP coupling, per night | 437.17<br>(255.33 – 605.17) | 521.83<br>(342 – 730.33) | 490.75<br>(333.33 – 658.75) |
| Low transition frequency SW-TB coupling | 2,836<br>(1,752 – 3,629.50) | 2,996<br>(2,081 – 3,575) | 2,305<br>(1,883.50 – 3,491) |
| Low transition frequency SW-TB coupling, per night | 479<br>(285.83 – 631) | 531.17<br>(369.20 – 609.67) | 443.20<br>(353.57 – 630.67) |
| Low transition frequency SW-SP coupling | 4,448<br>(2,975 – 5,701) | 4,991<br>(3,347 – 5,811) | 3,847<br>(2,968 – 6,413) |
| Low transition frequency SW-SP coupling, per night | 753.17<br>(496 – 968.67) | 858<br>(627.50 – 986) | 803.17<br>(540.25 – 1,068.83) |

### Event-Matched Time-Frequency Spectrograms of Theta Bursts

Theta burst coupling to both high and low transition frequency SWs was appreciable with a distinct TB spectral event in averaged time-frequency spectrograms (Figure 2). The normalized (SW coupling specific) TB power was quantified from a TF-window (region of time-frequency space) surrounding the TB spectral event for each individual and averaged to make group comparisons after controlling for co-variables and multiple comparisons (Table 3). Quantifying the normalized EEG power of TBs nested with high transition frequency SWs demonstrated ∼ 6.56% less normalized power among the Aβ+CI group, compared to the Aβ-CU [95% CI: (−2.83, -10.15%), adjusted *p* < 0.001]. The high transition frequency SW-coupled normalized TB power was also ∼6.14% lower comparing Aβ+CI to Aβ+CU groups [95% CI: (−2.62, -9.54%), adjusted *p* < 0.001].

**Table 3.** Precision of coupling and TF-window EEG power are expressed as a percent difference between groups, adjusted for covariates, and controlled for multiple comparisons with via Tukey-Kramer adjustment. Correlations between precision of coupling and biomarkers are adjusted for covariates only. Abbreviations: Aβ = amyloid; CU = cognitively unimpaired; CI = cognitively impaired; IQR= interquartile range; TF= time-frequency; SW= slow wave; SP= spindle TB=theta burst; * = statistically significant when controlled for a false discovery rate (FDR) of 0.05.

|  | Perfect Difference Estimate | Adjusted Confidence Interval | Adjusted p-value |
| --- | --- | --- | --- |
| **Aβ+CU vs Aβ-CU** |  |  |  |
| High transition frequency SW-TB precision | -2.568 | -4.993 – -0.081 | **0.041** |
| High transition frequency SW-SP precision | -3.042 | -7.815 – 1.979 | 0.319 |
| Low transition frequency SW-TB precision | -1.938 | -4.126 – 0.301 | 0.104 |
| Low transition frequency SW-SP precision | -5.099 | -8.770 – -1.280 | **0.006** |
| High transition frequency SW-TB TF-window | -0.448 | -3.710 – 2.924 | 0.946 |
| High transition frequency SW-SP TF-window | 1.395 | -2.256 – 5.182 | 0.645 |
| Low transition frequency SW-TB TF-window | -0.097 | -3.497 – 3.423 | 0.998 |
| Low transition frequency SW-SP TF-window | 0.848 | -2.754 – 4.583 | 0.847 |
| **Aβ+CI vs Aβ-CU** |  |  |  |
| High transition frequency SW-TB precision | -1.707 | -5.336 – 2.060 | 0.517 |
| High transition frequency SW-SP precision | -4.676 | -10.916 – 2.002 | 0.213 |
| Low transition frequency SW-TB precision | -2.032 | -4.814 – 0.832 | 0.210 |
| Low transition frequency SW-SP precision | -4.853 | -10.896 – 1.600 | 0.170 |
| High transition frequency SW-TB TF-window | -6.561 | -10.146 – -2.834 | **<0.001** |
| High transition frequency SW-SP TF-window | -1.083 | -5.364 – 3.393 | 0.826 |
| Low transition frequency SW-TB TF-window | -6.446 | -10.471 – -2.241 | **0.002** |
| Low transition frequency SW-SP TF-window | -0.393 | -4.728 – 4.139 | 0.976 |
| **Aβ+CI vs Aβ+CU** |  |  |  |
| High transition frequency SW-TB precision | 0.883 | -2.742 – 4.644 | 0.832 |
| High transition frequency SW-SP precision | -1.685 | -8.104 – 5.182 | 0.818 |
| Low transition frequency SW-TB precision | -0.096 | -2.886 – 2.774 | 0.996 |
| Low transition frequency SW-SP precision | 0.259 | -6.016 – 6.953 | 0.995 |
| High transition frequency SW-TB TF-window | -6.141 | -9.537 – -2.617 | **<0.001** |
| High transition frequency SW-SP TF-window | -2.444 | -6.566 – 1.860 | 0.360 |
| Low transition frequency SW-TB TF-window | -6.356 | -10.175 – -2.374 | **0.001** |
| Low transition frequency SW-SP TF-window | -1.230 | -5.412 – 3.136 | 0.772 |
|  | **Correlation Estimate** | **95% Confidence Interval** | **p-value** |
| **CSF 42/40 & Coupling Precision** |  |  |  |
| High transition frequency SW-TB coupling | 0.197 | 0.060 – 0.327 | **0.005\*** |
| High transition frequency SW-SP coupling | 0.140 | 0.001 – 0.273 | 0.047 |
| Low transition frequency SW-TB coupling | 0.151 | 0.012 – 0.284 | 0.033 |
| Low transition frequency SW-SP coupling | 0.183 | 0.046 – 0.314 | **0.009\*** |
| **p-Tau & Coupling Precision** |  |  |  |
| High transition frequency SW-TB coupling | -0.201 | -0.330 – -0.064 | **0.004\*** |
| High transition frequency SW-SP coupling | -0.121 | -0.255 – 0.019 | 0.089 |
| Low transition frequency SW-TB coupling | -0.091 | -0.226 – 0.049 | 0.201 |
| Low transition frequency SW-SP coupling | -0.134 | -0.267 – 0.005 | 0.059 |
| **total Tau & Coupling Precision** |  |  |  |
| High transition frequency SW-TB coupling | -0.175 | -0.306 – -0.037 | **0.013\*** |
| High transition frequency SW-SP coupling | -0.056 | -0.193 – 0.084 | 0.432 |
| Low transition frequency SW-TB coupling | -0.033 | -0.171 – 0.106 | 0.640 |
| Low transition frequency SW-SP coupling | -0.123 | -0.257 – 0.016 | 0.082 |
| **Combined SW-TB/SW-SP metric** |  |  |  |
| Precision correlation with Aβ<sub>42</sub>/Aβ<sub>40</sub> | 0.273 | 0.132 – 0.390 | **<0.001\*** |
| Precision correlation with p-tau | -0.21 | -0.073 – 0.339 | **0.003\*** |
| Precision correlation with total-tau | -0.182 | -0.044 – -0.313 | **0.01\*** |

**Figure 2:**
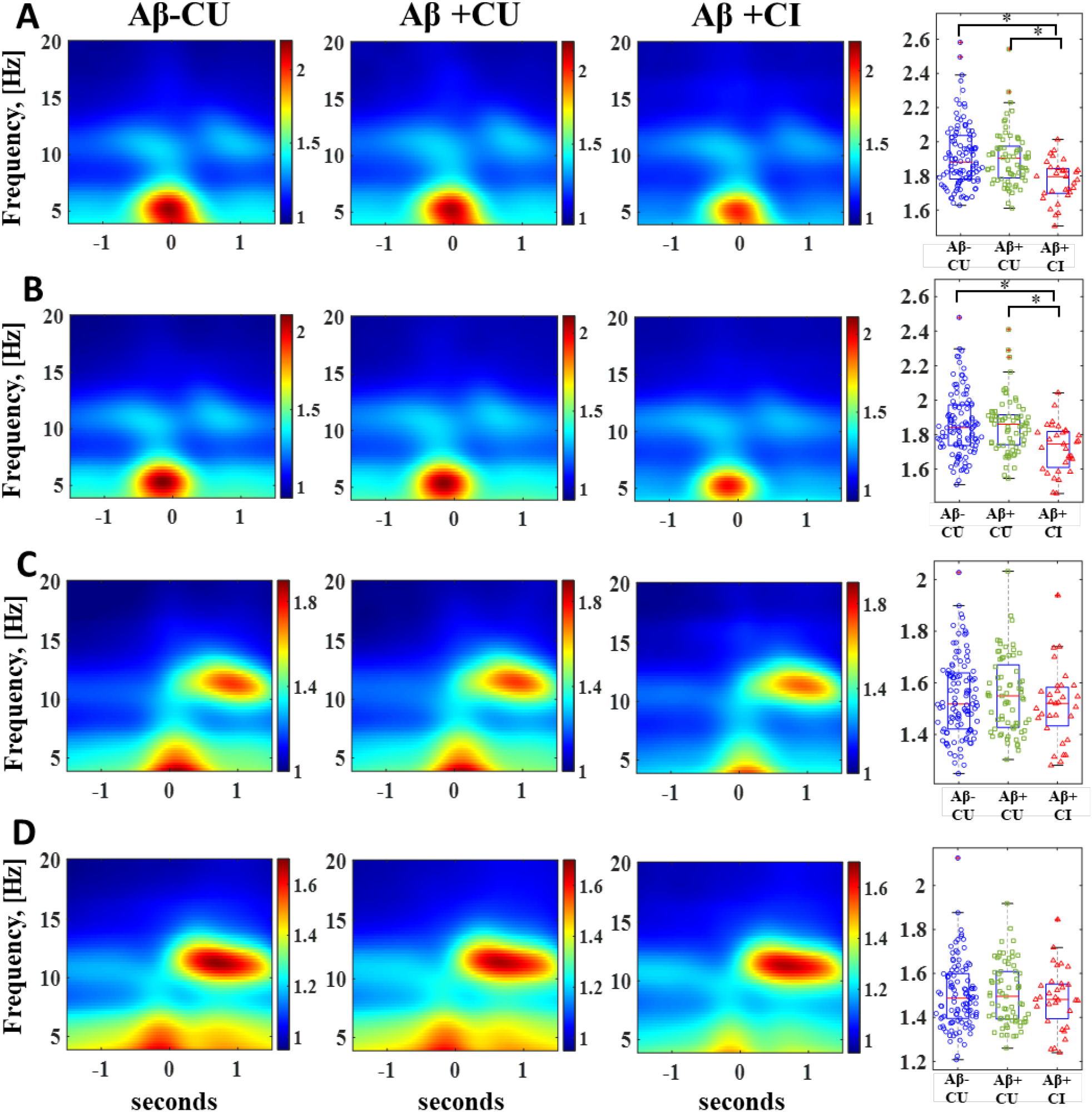
Comparison of SW-TB and SW-SP coupling normalized EEG power across stratified groups spanning stages of normal aging to mild cognitive impairment in early Alzheimer’s disease. A) High transition frequency SW-TB coupling and B) low transition frequency SW-TB coupling are visually represented as averaged time-frequency plots, and quantified EEG power among individuals is graphed in box/whisker plots. C) High transition frequency SW-SP coupling and D) low transition frequency SW-SP coupling are visually represented as averaged time-frequency plots, and quantified EEG power among individuals is graphed in box/whisker plots. Abbreviations: Aβ = amyloid, CU = cognitively unimpaired, CI = cognitively impaired, SP = spindle, SW = slow wave, TB = theta bursts. [*] indicates statistical significance <0.005 after adjusting for covariates and multiple comparisons (Tukey-Kramer adjustment).

The TF-windows for TBs matched to wide low transition frequency SWs demonstrated similar group comparisons, with ∼6.45% less EEG power among the Aβ+CI group compared to the Aβ-CU group [95% CI: (−2.24, -10.47%), adjusted *p* < 0.002]. The Aβ+CI group was ∼6.36% lower in TB power compared to the Aβ+CU group as well [95% CI: (−2.37, -10.18%), adjusted *p* = 0.001].

### Event-Matched Time-Frequency Spectrograms of Spindles

Time-frequency spectrograms for SPs matched to both high and low transition frequency SWs demonstrate a clear SP spectral event (Figure 2). As with TBs, normalized (SW-coupling specific) SP EEG power was quantified from a TF-window surrounding the SP spectral event for each individual and averaged to make group comparisons after controlling for co-variables and multiple comparisons (Table 3). Comparisons between groups demonstrated no statistically significant differences in SP TF-window normalized EEG values for either high or low transition frequency SWs.

### Event-Matched Precise Topographical Theta Burst Localization

The precision of SW-TB coupling for high transition frequency SWs demonstrated a modest, but statistically significantly ∼2.57% lower precision between Aβ+CU individuals compared to Aβ-CU individuals [95% CI: (−0.08, -4.99%), adjusted *p*<0.05] (Table 3). Additional group comparisons of precision of SW-TB coupling did not reach statistical significance after controlling for multiple comparisons (Table 3; Figure 3).

**Figure 3:**
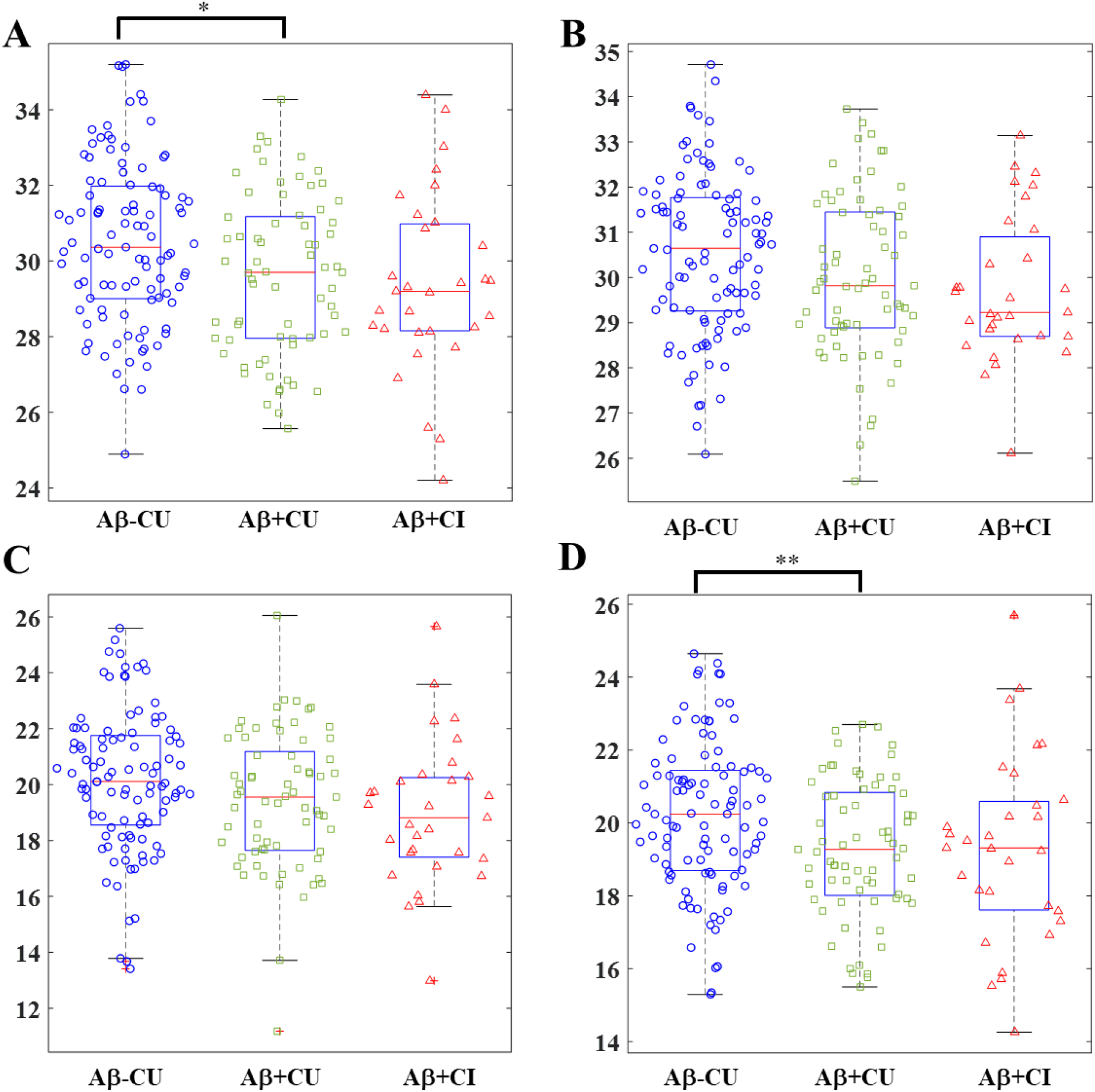
Comparison of SW-TB and SW-SP coupling precision across stages of normal aging to mild cognitive impairment in early Alzheimer’s disease. A) High transition frequency SW-TB precision comparison between groups. B) Low transition frequency SW-TB precision comparison between groups. C) High transition frequency SW-SP precision comparison between groups. D) Low transition frequency SW-SP precision comparison between groups. Abbreviations: Aβ = amyloid, CU = cognitively unimpaired, CI = cognitively impaired, SP = spindle, SW = slow wave, TB = theta bursts. [*] indicates statistical significance <0.05 and [**] indicates statistical significance <0.01 after adjusting for covariates and multiple comparisons (Tukey-Kramer adjustment).

### Event-Matched Precise Topographical Spindle Localization

The low transition frequency SW-SP coupling demonstrated ∼5.10% lower precision between Aβ+CU individuals compared to Aβ-CU individuals [95% CI: (−1.28, - 8.77%), adjusted p<0.01] (Table 3). Additional comparisons of precision within high and low transition frequency SW-SP coupling were not statistically significant between groups after controlling for multiple comparisons (Table 3; Figure 3)

### Correlations with CSF AD Biomarker Levels

We next performed an analysis of the relationships between SW-TB and SW-SP precision with concentrations of CSF biomarkers, controlling for age, sex, education, APOE4 gene status, AHI, and false discovery rate for multiple comparisons (Table 3). High transition frequency SW-TB coupling precision and low transition frequency SW-SP coupling precision were significantly correlated with CSF Aβ_42_/Aβ_40_ ratios (rho=0.197, 95% CI: (0.060 – 0.327), p=0.005; rho=0.183, 95% CI: (0.046 – 0.314), p<0.01). Neither low transition frequency SW-TB or high transition frequency SW-SP coupling precision were significantly correlated with Aβ_42_/Aβ_40_ levels. With regard to CSF p-tau and total-tau, high transition frequency SW-TB precision was the only coupling metric significantly correlated with CSF p-tau levels (rho=-0.201, 95% CI: (−0.330 – - 0.064), p<0.005) and with CSF total-tau levels (rho=-0.175, 95% CI: (−0.306 – -0.037), p<0.05). The correlations remained significant under a false discovery rate (alpha = 0.05) for all SW-TB and SW-SP precisions with Aβ_42_/Aβ_40,_ p-tau, and total-tau.

Given the significant AD biomarker correlations observed with high transition frequency SW-TB precision and low transition frequency SW-SP precision, we performed an additional exploratory analysis to observe whether combining these two distinct metrics might provide a better gauge of memory-associated neural circuit integrity. Here we combined these SW-TB and SW-SP metrics into one summed precision value (log scale precision values were summed for each participant to create a hybrid precision metric that incorporates both their SW-TB and SW-SP precision; Table 3; Figure 4). The covariate-adjusted correlations with this hybrid metric were statistically significant for CSF Aβ_42_/Aβ_40_ ratios (rho=0.266, 95% CI: (0.132, 0.390), p<0.001), p-tau (rho=-0.210, 95% CI: (−0.339, -0.073), p<0.003), and total-tau (rho=-0.182, 95% CI: (−0.313, -0.044)) p<0.010). The correlations remained significant under a false discovery rate (alpha = 0.05) for these summed precision values with Aβ_42_/Aβ_40,_ p-tau, and total-tau.

**Figure 4:**
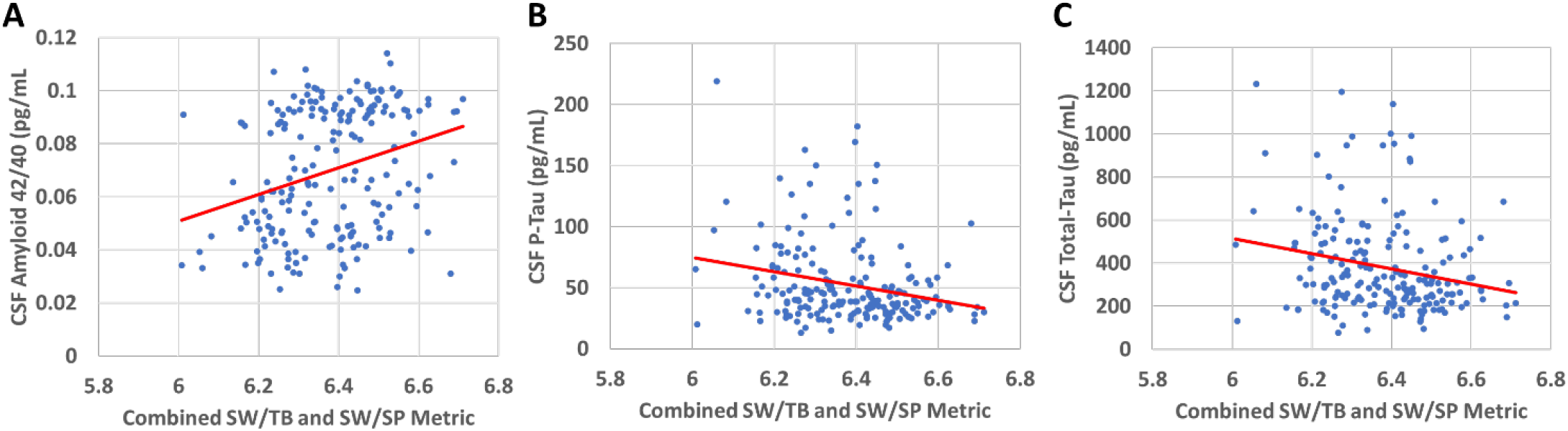
Regression analyses are illustrated for combined SW-TB and SW-SP metrics (log scale precision values were summed for each participant to create a hybrid precision metric that incorporates both their high frequency SW-TB and low frequency SW-SP precision). The covariate-adjusted correlations with this hybrid metric were statistically significant for A) CSF Aβ_42_/Aβ_40_ ratios (rho=0.266, 95% CI: (0.132, 0.390), p<0.001), B) p-tau (rho=-0.210, 95% CI: (−0.339, -0.073), p<0.003), and C) total-tau (rho=-0.182, 95% CI: (−0.313, -0.044)) p<0.010). The correlations remained significant under a false discovery rate (alpha = 0.05) for these summed precision values with Aβ_42_/Aβ_40,_ p-tau, and total-tau. Note that raw values are graphed for conceptual and illustrative purposes, while correlation coefficients and p-values were obtained via partial Spearman correlations after adjusting for covariate effects. Abbreviations: Aβ = amyloid, SP = spindle, SW = slow wave, TB = theta bursts.

## Discussion

Our study examined the properties of key brain communication events that are associated with memory replay sequences during slow wave sleep as an assessment of the potential biomarker properties of single channel sleep EEG in early stages of AD. Our results demonstrate distinctions between high and low transition frequency subtypes of SW events and the precision of neural circuits that control coupling of SWs to both TBs and SPs. Analysis of normalized SW-coupled TB EEG power among Aβ+CI individuals demonstrated significant differences in TB power coupled to both high and low transition frequency SWs, compared to both Aβ-CU and Aβ+CU individuals.

Amyloid positivity and ratios of CSF Aβ_42_/Aβ_40_ were associated with loss of precision in the circuits controlling high transition frequency SW-TB coupling and low transition frequency SW-SP coupling. Loss of high frequency SW-TB precision was also correlated with CSF p-tau and total-tau levels (see Figure 5 for an illustrative summary). An exploratory analysis further enhanced these correlative relationships by combining the high transition frequency SW-TB and low transition frequency SW-SP metrics, suggesting that these separate neural circuit metrics jointly contribute to each individual’s statistical relationship with AD core biomarkers. Taken together, our analyses reveal distinct abnormalities in memory-associated oscillatory events of SWA in association with both CSF biomarkers of Alzheimer’s disease and clinical symptoms of mild cognitive impairment as measured by CDR.

**Figure 5:**
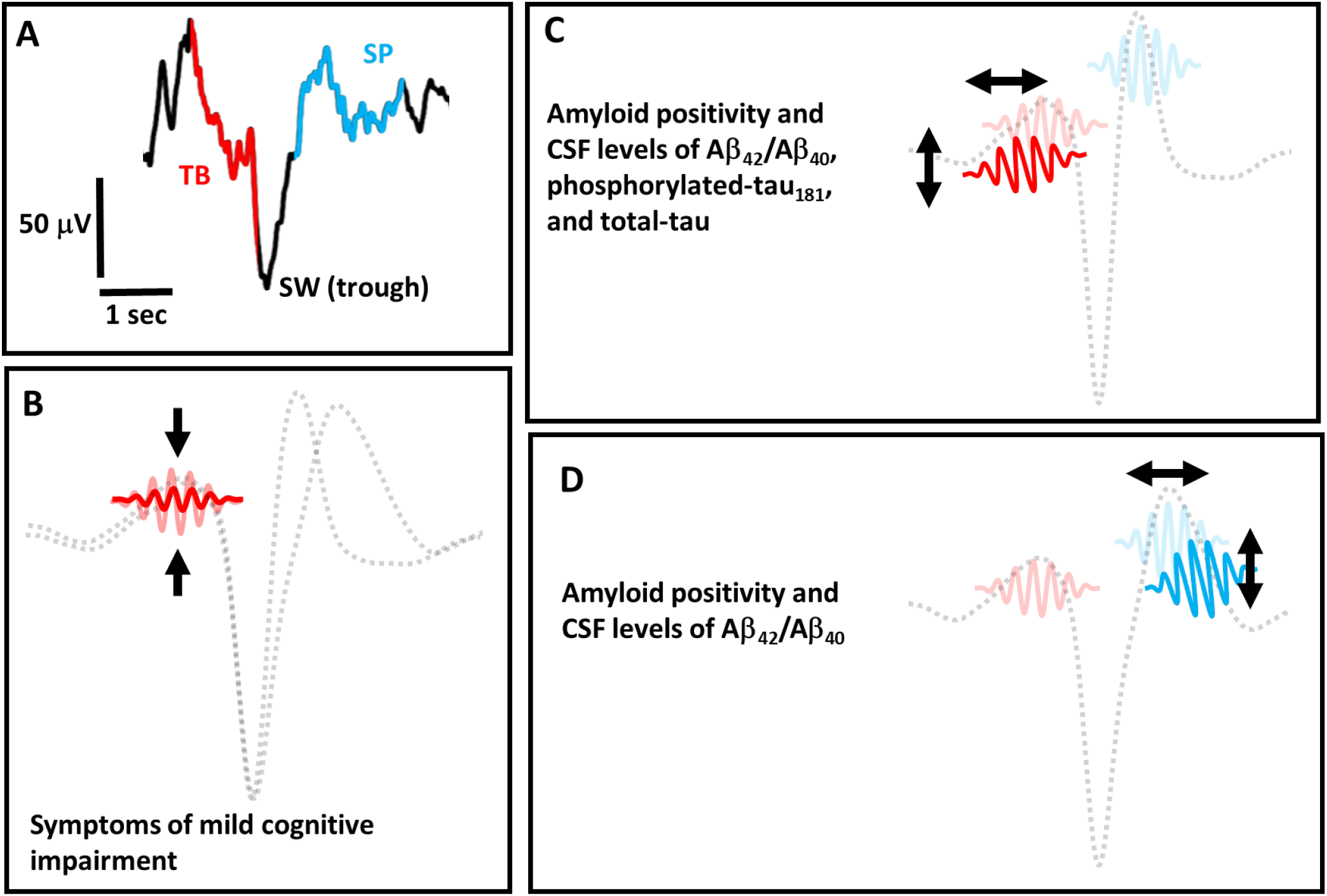
A) The temporal coupling of a single theta burst (TB) event and a single spindle (SP) event to the trough of a slow wave (SW) event is re-illustrated from Figure 1 as a conceptual reference. B) An illustration of reduced EEG power in TB events that are coupled to both high transition frequency and low transition frequency slow wave (SW) events (illustration not to scale). Individuals with symptoms of mild cognitive impairment by CDR testing exhibit relatively lower EEG power of TB events. C) A conceptual illustration of TB temporal and frequency precision “drift” in the coupling of TB events to high transition frequency SW events (illustration not to scale). Lower precision in this SW-TB circuit is associated with categorical amyloid positivity by CSF threshold levels, as well as higher CSF levels of Aβ_42_/Aβ_40_, phosphorylated-tau_181_, and total-tau. D) A conceptual illustration of SP temporal and frequency precision “drift” in the coupling of SP events to low transition frequency SW events (illustration not to scale). Lower precision in this SW-SP circuit is associated with categorical amyloid positivity by CSF threshold levels, as well as higher CSF levels of Aβ_42_/Aβ_40_. Abbreviations: Aβ = amyloid, CDR = Clinical Dementia Rating Scale. CSF = Cerebrospinal Fluid, SP = spindle, SW = slow wave, TB = theta bursts.

Our analyses focused on subtypes of SWs, defined by their transition frequencies, as well as TBs and a lower frequency (late-fast) subtype of SPs (higher frequency, early-fast spindles were not reliably measured from the single frontal channel). Notably, there were no significant differences in the average number of detected events between participant groups, nor were there significant differences in the number of detected coupled event pairs between participant groups. Sleep staging further demonstrated no significant differences between groups. Together, this suggests that that the production of the SWA-associated oscillatory events examined herein is not significantly impacted by amyloid positivity and/or symptoms of mild cognitive impairment, and instead implicates imprecision in the timing and frequency characteristics of these oscillatory events as early signs of neurodegenerative change.

Distinct oscillatory events found within SWA are generated in conjunction with replay of memory sequences, mirroring wake-like experiences in the patterns of neuronal activity^17-23^. Our work builds on previous reports that have described changes in these oscillatory events in conjunction with specific neuropathologies, including age-related atrophy and Alzheimer’s disease pathology^4,8,13,33,34,45-47^. Notably, measurement of oscillatory event coupling in these foundational studies relied on simple timing metrics, while our analyses deployed novel signal processing methods to calculate both time and frequency “drift” in the underlying SW-SP (and SW-TB) neural circuits for both high and low transition frequency SW coupling. Our study represents, to the best of our knowledge, the first formal description of this technique in sleep EEG analysis and is also the first study to expand coupling analyses to include SW-TB relationships.

The rationale for our approach rests on previously reported correlations between tau^34^ and amyloid^33^ PET positivity with SW-SP timing and on advances in signal processing that differentiated high versus low transition frequency SW-SP coupling in the context of aging^35^, amyloid positivity^33^, and cognitive impairment^33,36^. Our analyses corroborate these distinctions between high versus low transition frequency SW coupling precision both in association with CSF threshold-based amyloid positivity and in correlative relationships with Aβ_42_/Aβ_40_, p-tau and total tau, suggesting that SW-TB and SW-SP coupling differ in high versus low transition frequency categories. Conversely, utilizing a different metric of normalized EEG power in both high and low transition frequency SW-TB coupling resulted in similar distinctions among the individuals with mild cognitive impairment, while normalized EEG power for SW-SP coupling did not differ in group comparisons. These results suggest that measuring normalized EEG power versus the precision of event coupling quantify different neurophysiological processes.

Distinctions between SW-TB and SW-SP coupling are apparent in intracranial recording analyses, wherein the TB events are observed to initiate SWs and precede hippocampal sharp wave ripples, while SPs seem to coordinate high frequency gamma activity following the TB and sharp wave ripple events^25,26,37^. Notably, in the wake state, theta oscillations and sharp wave ripples are mechanistically linked to hippocampal pattern completion and reinstatement of memory tracings during recall of memory^48,49^.

Further, theta oscillations appear to play a similar role in both wake and sleep in coordinating the reactivation of recently learned memory sequences^38^. While speculative, it is possible that the same memory-reinstating theta circuits of wake-state are responsible for producing TBs during memory replay cycles within NREM sleep.

This connection between wake-state memory performance and sleep-state memory processing may explain why aging adults within the Aβ+CI group are observed to have lower theta burst power in their sleep EEG. Notably, SP precision abnormalities were not statistically different in comparisons between the Aβ+CI group with either Aβ-CU or Aβ+CU individuals, although the relatively lower number of CDR positive participants and high variance among this group may have contributed to lack of statistical significance.

While the dataset utilized for our analyses is significantly larger than prior studies of SW-SP coupling and Alzheimer’s-related pathology^33,34^, limitations in sample size may obscure possible differences between groups in the number of identified oscillatory events and sleep stage metrics, particularly in comparisons with the Aβ+CI group.

Further, although our utilization of a single channel of at-home EEG is highly advantageous for future translation of this method to inexpensive “wearable” devices, there are potential limitations in the use of an EEG headband device. Most notable among these technical constraints is the inability to reliably measure a category of “early-fast” spindles, as these higher frequency spindles are more prominent in posterior recording sites.^5^ In addition, the cross-sectional nature of this study limits our ability to examine potential longitudinal relationships between our novel sleep EEG metrics and neurodegenerative processes, and future studies will be required to explore potential predictive properties of sleep EEG as a biomarker.

Detection of oscillatory events in sleep may provide a clinical application as a marker of brain health and in early detection of neurodegenerative processes. This signal processing technique requires only a single channel of EEG that can be recorded in an unsupervised home setting, thus opening the door for development of brain health monitoring via relatively inexpensive and easily self-applied EEG “wearable” headbands. This technique is also relatively cost effective for potential longitudinal monitoring of both neurodegenerative disease risk and response to interventional treatments. Further, the metrics from sleep EEG may provide functional information related to memory circuits and is not susceptible to learning effects or volitional aspects of cognitive testing. Nonetheless, significant work remains to translate this technique to clinical application, including the need to refine the signal processing technique and more fully account for variance among individuals. Advancements in event detection and subtyping may provide a means to increase the precision of this technique. Considerable work also remains to determine the potential causal relationships between EEG events and neuropathology. Future studies will be required to catalogue dynamic changes in the EEG signals in response to improvement or worsening of neurophysiological processes that impact sleep’s memory processing system.

In conclusion, our data demonstrate that the time-frequency spectral properties of both TB and SP coupling to SWs can be precisely measured from single-channel EEG and provide information about the integrity of neural circuits controlling sleep’s memory replay in the early stages of AD pathogenesis. Changes in the integrity of these hippocampal-dependent memory circuits occur prior to development of cognitive symptoms and may serve as an early biomarker of neurodegenerative processes. Cross-sectional correlations with CSF AD biomarkers further suggest that SW-TB and SW-SP coupling are important processes to consider in the search to identify fundamental neuroprotective properties of sleep that may serve as targets for novel therapeutic development.

## Acknowledgements

The authors are grateful for data and resources provided by the Knight Alzheimer Disease Research Center and for resources provided by the University of Colorado Alzheimer’s and Cognition Center.

## Funding

This study was supported by the following grants from the National Institutes of Health: R01AG058772, P01AG003991

## Competing Interests

R.L.P. is employed by Google; V.O.K. consults for Garmin; D.M.H. co-founded and is on the scientific advisory board of C2N Diagnostics. D.M.H. consults for Genentech, Denali., Cajal Neurosciences, and Alector. Washington University receives research grants to the laboratory of D.M.H. from C2N Diagnostics and NextCure. J.C.M. is funded by NIH grants P30 AG066444; P01AG003991; P01AG026276 and U19 AG032438. Neither J.C.M. nor his family owns stock or has equity interest (outside of mutual funds or other externally directed accounts) in any pharmaceutical or biotechnology company. B.P.L. consults for Eli Lilly, has funding from Eisai, and is a member of the scientific advisory board for Beacon Biosignals; E.K., L.M.M., A.R.R., C.D.T., S.H.S., B.M.B., and B.V.M., declare that they have no competing interests.

